# Cold stress involves CBF dependent regulatory pathway to remodel wood formation in *Eucalyptus gunnii* hybrids

**DOI:** 10.1101/2024.12.11.627904

**Authors:** Ines Hadj Bachir, Raphael Ployet, Hélène San Clemente, Marielle Aguilar, Annabelle Dupas, Nathalie Ladouce, Yves Martinez, Jacqueline Grima-Pettenati, Chantal Teulières, Hua Cassan-Wang, Fabien Mounet

**Author notes:** Author for correspondence: Mounet Fabien.

## Abstract

While being the most planted tree worldwide, most *Eucalyptus* species are sensitive to frost. Long-term exposure to cold temperatures, among other abiotic cues, triggers modification of secondary xylem differentiation in *Eucalyptus*, but the molecular mechanisms behind it remain unknown. Overexpression of key players of cold signalling pathway, the CRT-repeat binding factors (CBF), not only causes the expected increase in freezing tolerance but also remodels wood in a similar trend to cold acclimation, making it a good candidate for wood functional adaptation to cold stress. To gain insight in CBF role in cold-induced secondary cell wall (SCW) remodelling, we used both targeted and untargeted methodologies to show that chilling and freezing temperatures induced the deposition of a thick SCW with alterations in lignin and polysaccharides composition as well as modifications of wood anatomy in a *Eucalyptus* cold-tolerant hybrid. Using co-expression network approaches, we identified CBF transcription factors (TFs) as a regulatory hub in xylem cold response. Direct targets of the CBF TFs were identified by DAPseq and unravelled promising candidates involved in SCW deposition and hormonal signalling pathways. Our results shed new light on the interplay between cold response and wood formation, bringing new evidence for the role of the cell wall in trees tolerance to abiotic stresses.

## 1. Introduction

In the current context of global warming, extreme weather events such as drought and heat waves are more likely to occur and threaten tree species survivability (Tomé et al., 2021). Besides the global increase of mean surface temperature, a drier climate is predicted within many European areas (IPPC, 2014), which would likely deeply modify the dynamics and repartition of both natural and cultivated species. The use of highly productive tropical trees like eucalyptus is important for wood production in Europe and represents a promising alternative to face climate change (Booth, 2013). Due to its inherent qualities for industrial plantations (high productivity, short rotation time and wide adaptability), *Eucalyptus* is the most planted hardwood species and one of the best sources of wood biomass worldwide (Booth, 2013; Fernández et al., 2018). Eucalyptus is economically important in Southern Europe and its cultivation area is supposed to move towards northern regions, at higher altitudes (Deus et al., 2018), where the most productive species *E.grandis* and *E.globulus*, particularly sensitive to winter frost, can hardly be planted. Eucalypts genus encompass >700 species originating almost exclusively from Australia and Tasmania (Brooker, 2000) and growing in a wide variety of conditions including high altitude land, like *Eucalyptus gunni* (Potts, 2004). Hybrids of *E.gunni* represent promising genotypes for forest plantations as well as an invaluable tool to study low temperature effects on wood developmental processes.

Wood, or secondary xylem, is the most abundant biomass on earth (Bar-On et al., 2018) and is formed by an internal meristem, the vascular cambium, through a differentiation process. Cambial cells divide, expand and a thick secondary cell wall (SCW) mainly composed of cellulose, hemicelluloses and lignin is deposited before undergoing programmed cell death (Plomion et al., 2001). This differentiation process gives rise to a complex and heterogeneous tissue made of different cell types, fulfilling the various physiological functions of wood: vessels for water and sap conduction, fibres for mechanical strength and ray cells for nutrient storage (Déjardin et al., 2010). Wood structure and composition are modified according to environmental cues and this plasticity is a way for perennial plants to adapt to abiotic stresses (Zinkgraf et al., 2017). In woody perennials, vessels and fibers size, xylem density, SCW thickness or composition are modified in response to drought, salt exposure, nutrients depletion or extreme temperatures (Hori et al., 2020; Ployet et al., 2019; Wildhagen et al., 2017; Yu et al., 2021).

Wood formation is tightly regulated in time and space by a hierarchized multi-layered network of transcription factors (TFs). The first tiers of this transcriptional regulatory network (TRN) include TFs from diverse families which regulate the expression of structural genes involved in xylogenesis and SCW deposition (Hussey et al., 2013; Nakano et al., 2015; Zhang et al., 2018; Ohtani and Demura, 2019). They are in turn controlled by tiers 2 and 3 master regulators, belonging mainly to MYB and NAC families. As recently reviewed by Hussey et al. (2022), a fourth and even a fifth tier of TFs, tightly linked to auxin hormonal signaling pathway, ensure an upstream regulation of tier 2 and 3 TFs. Genes of SCW TRN can be co-opted (i.e. diverted from their initial role in xylem development) by distinct abiotic stresses, to promote functional adaptation to abiotic constraints (Hadj Bachir et al., 2022). In *Arabidopsis*, high salinity and iron-deprivation were shown to alter the expression of SCW regulators (Taylor-Teeples et al., 2015). In woody perennials, the regulatory hubs interconnecting stress signalling pathways and wood regulatory network are poorly characterized but the few that were identified triggered modification of xylem formation. In birch, *Bp*NAC012, the closest ortholog of *At*SDN1, a master regulator of SCW TRN, is able to bind to the promoter of both SCW associated genes and abiotic stress responsive genes, enhancing salt and osmotic stress tolerance (Hu et al., 2019). *Egr*MYB137, a novel SCW TF acting at the crosstalk between wood formation and stress response in *Eucalyptus*, contributes to increase SCW thickness in xylem fibres (Ployet et al., 2019). *EgrMYB137*’s closest orthologs in poplar, *PtrMYB074* is involved in drought tolerance by regulating *Ptr*bHLH186 (Chen et al., 2019a; Liu et al., 2022). Most of the target genes of *Egr*MYB137 and *Ptr*MYB074 are related to SCW deposition and lignin biosynthesis (Liu et al., 2022; Takawira et al., 2023). In *Betula platyphylla*, *Bpl*MYB46, ortholog of *At*MYB46, a key regulator of SCW deposition (Ko et al., 2014), is also associated with enhanced salt and osmotic stress tolerance. BplMYB46 regulates key ROS-scavenging enzymes, enhances lignin deposition and SCW thickness (Guo et al., 2017). Similarly, in apple trees (*Malus domestica*), *Md*MYB46 not only binds to the promoters of lignin biosynthetic genes but also stress-responsive genes belonging to the AP2/ERF (APETALA2/ETHYLENE RESPONSE FACTOR) family, leading to an increased lignin deposition coupled with better abiotic stress tolerance (Chen et al., 2019b).

AP2/ERF TFs can regulate genes belonging to the SCW TRN to promote functional adaptation to various stresses. In *Arabidopsis*, rice and carrot, ERF TFs were shown to increase lignification while promoting drought tolerance (Ambavaram et al., 2011; Jung et al., 2021; Lee et al., 2016; Li et al., 2020). In Populus, ERF139 was described as a transcriptional regulator of xylem cells expansion and SCW formation, possibly in response to osmotic adjustments (Wessels et al., 2019). Overexpression of CBF (C-repeat binding factors), belonging to DREB1 group of AP2/ERF (Cao et al, 2015), triggers the deposition of a thicker and more lignified SCW, a reduction of cambial cell layers, a decrease of vessel size and an increase of vessel density in transgenic *Eucalyptus* (Cao et al., 2020). CBF TFs, which are major regulators of cold signalling pathway in plants, can bind conserved CRT/DRE *cis* elements within target genes promoters to induce their expression, thus contributing to the plant’s cold acclimation process (Park et al., 2015; Stockinger et al., 1997). In young and adult *E. gunnii* hybrids submitted to chilling temperatures, CBF genes were induced in xylem together with SCW-related genes, SCW was thicker, lignin content was increased and S/G ratio was decreased in xylem tissues (Ployet et al., 2018). Despite several evidence that cold signalling influences wood formation as part of the acclimation process, no evidence was brought that CBF can directly regulate genes involved in wood-related TRN.

In the present study, we aimed at deciphering the interconnected gene network between CBF-mediated cold signalling pathway and wood formation process in *Eucalyptus*. Taking advantage of an untargeted and large scale transcriptomic analysis coupled to wood anatomy and composition phenotyping approaches, we identified transcriptionally coordinated modules of genes potentially involved in the modification of wood properties induced by low temperature. We combined DAPseq approach with transcriptomic data to understand how *DREB1/CBF* genes could modulate the wood differentiation process during chilling and freezing stresses.

## 2. Materials and methods

### 2.1. Plant material and sampling

6-month old *Eucalyptus gundal* clones (*E.gunnii* x *E.dalrympleana* hybrids) provided by the FCBA (Pierroton, France) were submitted to a cold acclimation and a freezing stress. More than 150 clonal copies of *E.gundal* were grown in a growth chamber at 25°C/22°C (16 h light/ 8h dark) and then submitted to a long term cold exposure at 4°C during 46 days, followed by a freezing stress at -5°C for 2 days, before one week of recovery at 25°C. The most lignified part of the stems was harvested at 25°C before temperature decrease, after 2 days and 46 days at 4°C, after 48h at 4°C/-5°C (day/night) and after one week of recovery at 25°C (Figure S1). For histological analysis, a 1cm cylinder of each stem was kept apart in 80% ethanol. The remaining stems were debarked, cut in 0.5cm cylinders, frozen in liquid nitrogen. For each harvesting point, 18 plants were sampled to obtain 6 pools of xylem tissues from 3 independent plants. For transcriptomic and biochemical analysis, xylem samples were ground to powder using a Mixer Mill MM400 (Retsch) and kept at -80°C until further use.

### 2.2. Microscopy and histochemistry

For cell imaging and micro-phenotyping, stem sections were embedded in LR White resin (Electron Microscopy) by successive baths of increasing LR white concentration during 2 days (25%, 50%, 75%, 100%). Semi-thin sections (1 µm) were obtained using an ultramicrotome (Reichert Ultracut) and a Histo Diamond Knife (DiATOME). Sections were stained with Toluidine Blue O (0.1%) for 10 min before rinsing with deionized water. Images were subsequently acquired using a Nanozoomer and NPview2 software (Hamamatsu). Images were analysed with ImageJ software (Schneider et al., 2012) to measure various cell parameters: SCW thickness of fibres, vessel area, vessel density and number of cambial cell layers. SCW thickness was measured on the fibre cells within the first 3 or 4 cell layers below the cambium using TOASTER (ImageJ plugin) on approximately 200 fibre cells per biological replicate (6 biological replicates per time point). Vessels analysed were situated no more than 100 µm away from the cambium, corresponding to 70 to 220 analysed vessels per time point. To assess cambium activity, the number of cambial cell layers were counted on 5 distinct zones for each biological replicate (6 biological replicates per time point). Statistical significance was assessed by Student’s t-test.

For histochemical analyses of xylem tissues, cross sections of 60 µm and 200 µm were cut using a Vibratome (Leica VT 1000S). 60 µm cross sections were stained with Phloroglucinol-HCl and observed with an inverted bright-field microscope (Leica). 200 µm cross sections were fixed in glutaraldehyde 2% and 0.1 M Sorensen for 1h at 4°C. The samples were then dehydrated in successive ethanol baths of increasing concentration followed by a desiccation performed by a EM CP300 automated critical point dryer (Leica) with CO_2_ as a transitional fluid. When dry, the samples were coated with platinum (4 to 10 nm) in a EM MED020 vacuum coating system (Leica) and observed in a scanning electron microscope (ESEM Quanta 250 FEG FEI, Merignac, France) at 5 kV.

### 2.3. FT-IR analysis

Xylem samples were freeze-dried for 48h and extractives were removed by successive baths of boiling water, boiling 100% ethanol, boiling toluene/ethanol (50/50, v/v) and acetone. The resulting extractive-free xylem residues (EXR) were analysed by Fourier transformed infra-red spectroscopy (FT-IR) with an attenuated total reflection (ATR) Nicolet 6700 FT-IR spectrometer (Thermo Fisher) equipped with a deuterated-triglycine sulfate (DTGS) detector. 30 samples were analysed corresponding to 6 biological replicates per time point. Spectra were acquired between 4000 and 400 cm^-1^ with a 4 cm^-1^ resolution and 32 scans per spectrum. Median spectra were calculated from ten technical replicates for each sample. Baseline correction, normalization and offset correction was performed using R packages (hyperspectr, prospect and base respectively). Partial least square-Discriminant analysis (PLS-DA) was performed using mixOmics R package (Rohart et al., 2017).

### 2.4. RNA extraction and analyses of the transcriptome

RNA was extracted from ground xylem as described by Muoki et al., (2012) on the 30 samples described earlier. Contaminating DNA was removed using the TURBO DNA-free kit (Invitrogen). Six sequencing libraries per time point were generated and sequenced (Illumina, San Diego, CA, USA; 150 bp paired-end reads) by the Genotoul GeT platform (https://get.genotoul.fr/en/; France) using the HiSeq3000 (Illumina). An average of 40 million reads for each sample was obtained.

After curation with TrimGalore, reads were mapped on *E.gunnii* genome using HISAT. To perform differential analysis, htseq-counts files were analysed with EdgeR package version 3.24.3 (McCarthy et al., 2012). First, genes which did not have at least 1 read after a count per million normalization in at least one half of the samples, were discarded. Then, raw counts were normalized using TMM method and count distribution was modelled with a negative binomial generalized linear model where the time and the replicate were taken into account and where the dispersion is estimated by the edgeR method. Raw p-values were adjusted with the Benjamini–Hochberg procedure to control the False Discovery

Rate (FDR). A gene was detected as differentially expressed (DEG) if |log2(expression fold change) |>1 and FDR adjusted p-value < 0.05 between two time points of the kinetics. 8236 DEGs during the cold treatment were identified. Gene-gene correlations between the DEGs were calculated using Pearson pairwise correlation and the corresponding matrix was generated using the “rcorr” function of the “Hmisc” R package. The p-values associated to the correlation value were adjusted by the FDR correction method. Genes displaying correlations with a FDR-adjusted p-value < 10^-6^ were used to construct a co-expression network visualized with Cytoscape software, resulting in 8233 nodes linked by 4 278 476 correlations. Modules of genes were identified using the WGCNA method (Langfelder and Horvath, 2008) with the following characteristics: softPower=14, deepSplit=3 and minModuleSize=280. Gene Ontology enrichment analyses were performed on these modules using g:profiler (Raudvere et al., 2019). *Egu*CBFQ-subnetwork was obtained by extracting g58022 direct neighbours from complete network (correlations filtered with FDR-adjusted p-value <10^-6^), resulting in 1106 nodes linked by 515 620 edges. Gene ontology was performed using BiNGO from Cytoscape software (Maere et al., 2005).

### 2.5. DAPseq analysis

*EguCBFQ* (g58022, *E.gunnii* closest ortholog of *EgrCBF14*) coding sequence was isolated by PCR with corresponding primers (Table S1) from previous clones (Cao et al., 2020) obtained from *Eucalyptus gunnii* genomic DNA. *EguCBFQ* coding sequence was first cloned into pENTR/D/TOPO vector by BP reaction (Gateway BP Clonase II, Invitrogen), and then transferred to pIX HALO vector (CD3-1742, Tair) by recombination (Gateway LR clonase II, Invitrogen).

DAPseq experiment was performed following the protocol described in Bartlett et al., (2017) with minor optimization steps described in (Takawira et al., in prep). 5 µg of genomic DNA, extracted from xylem of *E.gundal* cold tolerant clone (named 208) harvested in June (optimal growth conditions) with the CTAB method, was sonicated at 200 bp using BioRuptor Pico (Diagenode). The DNA library was prepared thereafter: DNA was end-repaired and A-tailed using Klenow Fragment (3’-5’) enabling Illumina TruSeq adaptors ligation. In parallel, to obtain HALO-fused CBF proteins, the pIX-HALO vector containing CBF CDS was used for *in-vitro* transcription/translation using SP6 TnT® Quick Coupled Transcription/Translation System (Promega). The HALO-fused CBF proteins were purified thanks to Magne HaloTag beads (Promega). A control sample corresponding to HaloTag protein alone was used (referred to as pIX-HALO empty). Correct expression of the HALO-CBF protein was verified by Western Blot analysis with an anti-HALO primary antibody. The HALO-CBF TF or HaloTag alone were then incubated with 100ng of genomic DNA library. Bound DNA-fragments were amplified by PCR thanks to specific primers (Table S1) corresponding to Illumina adaptors. Amplified libraries were pooled together before size selection around 400bp with magnetic beads (SPRIselect, Beckman Coulter). The resulting DNA pool was used for further Illumina sequencing on Novaseq6000 to obtain 2x150 paired-end reads. Sequencing data treatment was performed according to the workflow established in Figure 4. Three independent pIX-HALO empty libraries were merged and used as a negative sample set for peak calling. Peak calling was performed using MACS2 and peaks were visualized using Integrative Genome Viewer (Robinson et al., 2011). A peak was assigned to a target gene if situated within 5kb before TSS and 3 kb after stop codon. Identification of consensus motifs from sequences of 500 bp around the maximum of the peak was performed using MEME-ChIP (Machanick and Bailey, 2011).

## 3. Results

### 3.1. Cold modifies wood structure and composition

To investigate the effect of chilling and freezing temperatures on xylem structure and SCW composition, we performed histological analyses on cross sections of stems from young eucalyptus hybrids that have been submitted to cold acclimation (4°C), followed by a freezing stress during 2 days (4°C / -5°C, days / night) and a recovery at 25°C. At the end of cold acclimation, fibre cells situated within the first 3 to 4 cell layers immediately below the cambium (corresponding to the xylem differentiation zone) displayed a thicker cell wall in samples submitted to cold compared to samples grown at 25°C (Figure 1A-B, E-F, G). In this zone, vessels were smaller in cold-treated plants compared to control plants (Figure 1H). Besides, cold-treated samples had a reduced number of cambial cell layers (Figure 1I). Histochemical staining with Phloroglucinol-HCl revealed an increase of cell wall lignification in developing xylem cells within the first 4 cell layers below the cambium. This observation was particularly pregnant for axial parenchyma cells surrounding vessels (Figure 1C-D).

**Figure 1:**
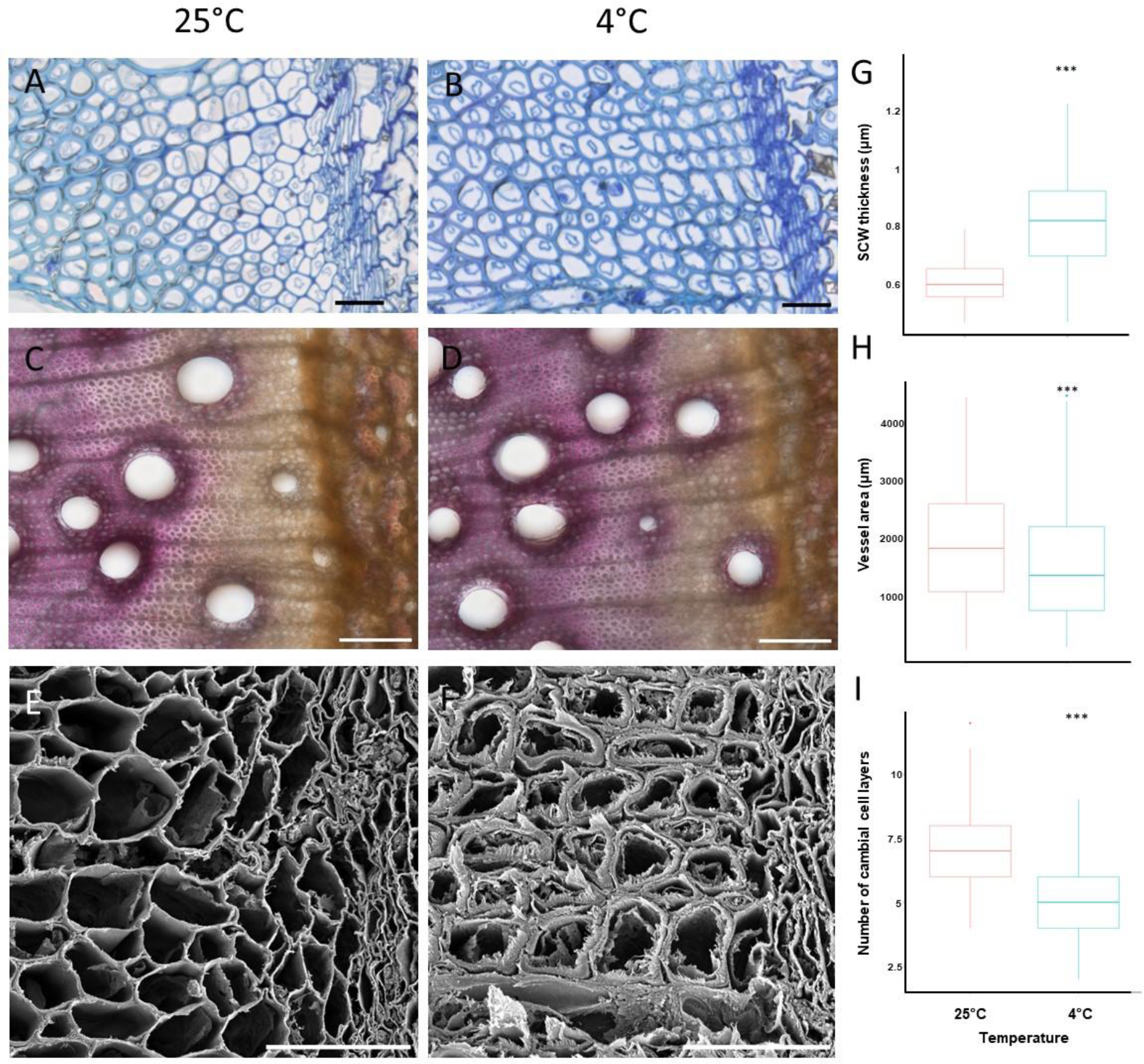
Cold triggers modifications of wood structure in cambial and developping xylem zones. **A to F**: Comparison of stem cross sections harvested from Eucalyptus grown at 25°C or 4°C (during 42 days). Tissues were stained with Toluidine blue (A and B) or Phloroglucinol-HCl (C and D). E, F: Magnification of cambial and developping xylem zones using scanning electron microscopy (x2000). Ca: cambium; P: phloem, mX: mature xylem, dX: differentiating xylem, X: xylem. Red arrows indicate lignifying vessels at 4°C. Scale bar: 20 µm (for A, B, E, F) and 100 µm (for C and D). **G to I**: Analyses of SCW thickness, vessel size and cambial activity in control (25°C) and cold acclimated (4°C) Eucalyptus plants. n= 6 to 18 independent plants. Image of Toluidine blue stained stem cross sections were acquired at 40x magnification using a Nanozoomer High Throughput (Hamamatsu) and NPview2 software (Hamamatsu) and analysed using ImageJ software (Schneider et al., 2012) and the TOASTER plugin. Asteriks indicate significant differences between control (25°C) and cold-treated samples (4°C): Student’s t-test: *p<0.05, **p<0.01, ***p<0.001

To further characterize global changes in SCW composition, we performed chemical fingerprinting analyses using Fourier transformed infra-red spectroscopy (FT-IR) on milled EXR samples before and after 46 days of cold exposure. A multivariate statistical analysis (Partial least-square discriminant analysis) applied on all the spectra obtained (from 1800 cm^-1^ to 800 cm^-1^) led to a clear separation between cold-treated and non-cold treated samples (Figure 2A). The first three components explained 77% of the total variance and PC1 is responsible for the separation between samples submitted to long-term cold exposure (4°C) and controls (25°C). This result indicates a global modification of SCW composition in response to cold. The analysis of the loadings contribution values to PC1 from the 500 most discriminant wavenumbers (WN) identified by Sparse-PLSDA combined to the comparison of FT-IR absorption spectra allowed the identification of wavenumbers explaining the separation between eucalyptus grown at 4°C and 25°C (Figure 2B). We identified 64 wavenumbers related to SCW components according to literature data accounting for the differences in EXR composition between cold exposed plants and plants grown at 25°C. We used loading values obtained from sparse PLS-DA to identify 6 zones of FT-IR spectra presenting the highest contribution to the separation between 4°C and 25°C conditions.

**Figure 2:**
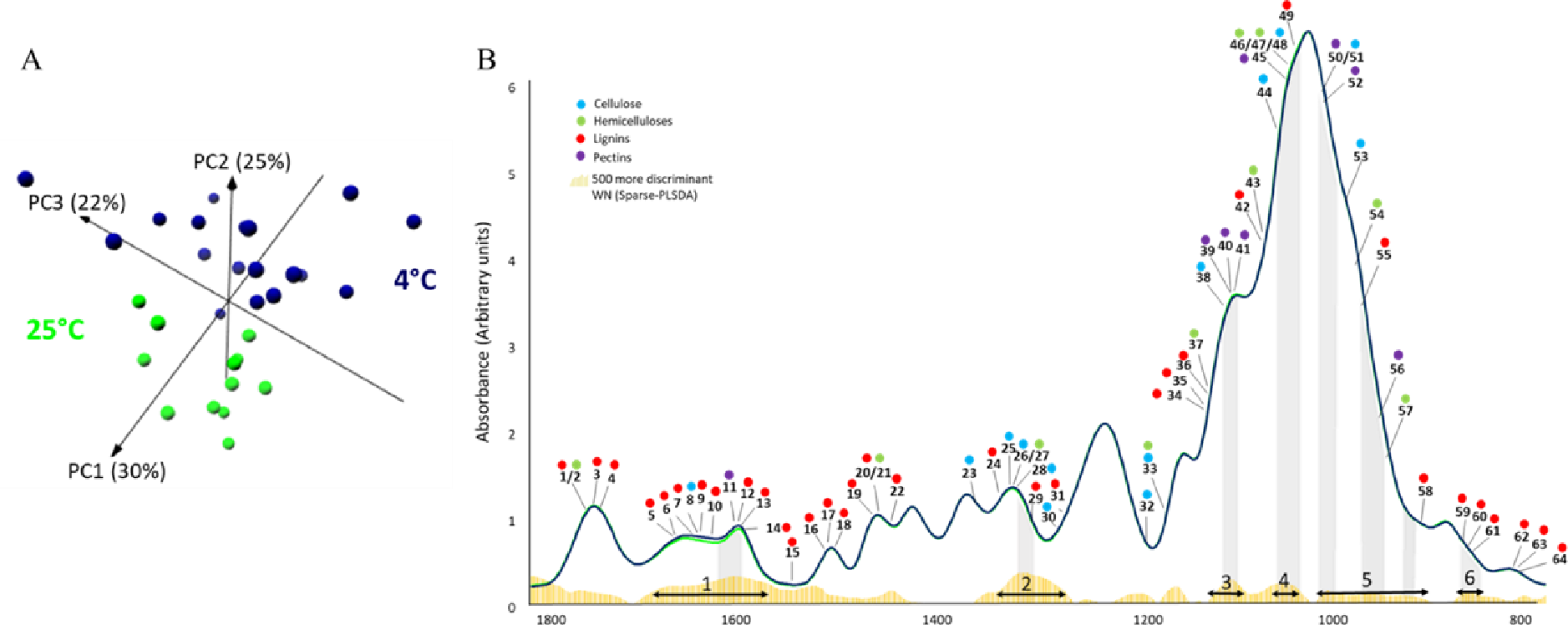
Global changes of SCW composition during cold acclimation revealed by FT-IR fingerprinting. **A:** 3D-Partial least square analysis (3D-PLS-DA) was performed using the normalized values of Fourier-Transformed Infra-Red (FT-IR) absorption spectra from 1800-800 cm^-1^. The first three components explain 77% of total variability and separates control samples (lightgreen dots) from cold-treated samples (darkblue dots). **B:** FT-IR median absorption spectra of control (lightgreen) and cold-treated samples (darkblue). Yellow histograms represent the absolute value of the contribution on PC1 of the 500 more discriminant wavenumbers (WN) identified by Sparse-PLS-DA. 64 FT-IR WN associated to cell wall compounds were identified according to literature data (blue, green, red and purple dots were associated to cellulose, hemicelluloses, lignins and pectins respectively. See Supplementary Table S2). Black arrows indicate WN zones responsible for most of the separation between samples and grey background highlights the contribution peak of each zone.

Zone 1 and zone 6 were mainly associated to lignin-related WN (7 out of 9 WN in zone 1 and 6 out of 6 WN in zone 6), whereas all the other zones were enriched in WN related to polysaccharides, mainly hemicelluloses and cellulose (Table S2). In zone 1, the absorption of cold-treated lines was clearly higher than in controls, indicating a putative increased lignification caused by cold treatment. Overall, these data highlight a global modification of SCW composition, and more precisely polysaccharides, induced by long-term cold exposure.

### 3.2. Chilling and freezing induce a reprogramming of transcriptional network in xylem cells

To gain an overview of the transcriptomic changes occurring in the stem of young eucalyptus during cold acclimation and freezing stress, we performed a RNAseq analysis on xylem samples. We identified 8236 differentially expressed genes (|log2(expression fold change)|>1 and FDR adjusted p-value < 0.05, n=6). We calculated Pearson correlations between the expression profiles of these DEGs and used them to build a co-expression network. We filtered the correlations according to their adjusted FDR p-value < 10 ^-6^ which resulted in a dense and highly correlated network of 8233 nodes linked by 4 278 476 edges (Figure 3). Using WGCNA and GO enrichment analyses, we detected 11 modules of co-expressed genes.

**Figure 3:**
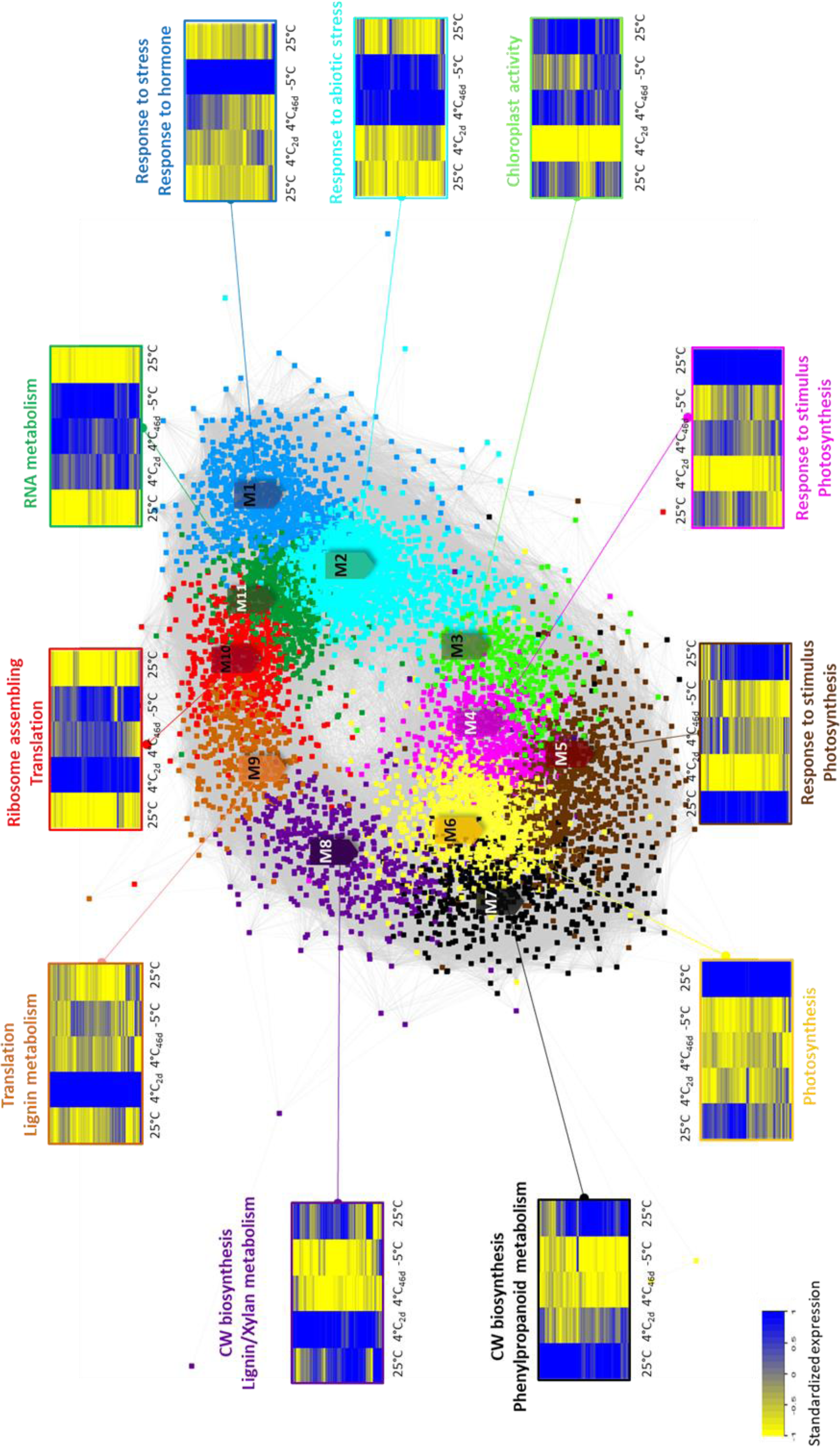
Identification of gene modules invovled in the trade-off between radial growth and cold tolerance of wood tissues by WGCNA. Weighted co-expression network was obtained by calculating Pearson correlation values between standardized expression profiles of 8236 DEGs. Gene-gene correlations with an adjusted-FDR p-value < 10^-6^ were used to generate the co-expression network, resulting in 8233 nodes linked by 4 278 476 edges, visualized with Cytoscape software (Force directed layout). Edge length is proportional to correlation. Eleven modules of genes represented by different node colors were detected by WGCNA approach (Langfelder and Horvath, 2008). For each module, gene expression profiles along the cold kinetics (Figure S1) were represented by heatmaps; 4°C_2d_: after 2 days at 4°C; 4°C_46d_: after 46 days at 4°C. Gene Ontology (GO) enrichment analysis was performed with g:profiler using orthology between *E.gunni* and *A.thaliana* to identify biological processes associated with each module.

Modules M3, M4, M5, M6, M7 and M8 include 3 891 DEGs and were globally repressed during the cold treatment. Inside this global repression pattern, various expression profiles can be distinguished: (1) genes that were repressed during the whole cold treatment (module M6); (2) genes that were repressed at each temperature change (module M5); (3) genes for which the acclimation process had enabled them to find their classical expression level (module M3); and finally (4) genes that were repressed only after a long cold treatment (modules M7 and M8). According to Gene Ontology analysis, the repressed modules were enriched in genes related to growth and cell metabolism. More precisely, we detected enrichment in biological processes related to primary metabolism and resources remobilization (M3 to M6), as well as cell wall biosynthesis (M7 and M8).

On the contrary, there were also modules of genes that were induced during the cold acclimation and freezing stress (M1, M2, M9, M10, M11). These modules were composed of 4 344 DEGs. Inside these modules gathering cold-up-regulated genes, we found different expression patterns: (1) genes induced only at the beginning of the cold treatment (M9), (2) during the whole cold treatment (M10, M11), (3) only after a long period of cold treatment (M1, M2). These modules are enriched in genes related to abiotic stress response (M1 and M2, encompassing numerous stress-responsive genes, as well as hormonal signalling pathways, and genes involved in secondary metabolism, such as phenylpropanoid biosynthesis) and related to cell metabolism and activity (transcription, translation, RNA metabolism in M9, M10 and M11).

As we aimed at understanding SCW-remodelling observed in response to cold, we then decided to focus specifically on wood and cell wall related genes.

As stated before, the two modules (M7 and M8) that were clearly related to CW biosynthesis belonged to repressed modules. The heatmaps of these modules revealed a down-regulation of these genes after 7 weeks of cold treatment and during the freezing stress. Most of these genes were still expressed at T1. In these 2 modules, we found 6 genes belonging to the lignin *bona fide* pathway (*EgrPAL9, EgrPAL3 EgrHCT5, EgrCCoAOMT2, EgrCOMT1* and *Egr4CL1*) corresponding to 5 early catalytic steps of lignin biosynthesis. There were also 14 genes involved in hemicelluloses biosynthesis and 6 cellulose biosynthesis genes. 6 known SCW regulators were clustered in these modules: two early master regulators *EgrNAC49* (ortholog of *AtNST1/2*) involved in fibre development and *EgrNAC26* (*AtVND6*) involved in vessel development, two master switches *EgrMYB2* (*AtMYB46*) and *EgrMYB31* (*AtMYB46*) as well as other MYB/NAC TFs, namely *EgrNAC152* (*AtXND1*) and *EgrMYB61* (*AtMYB85*). We also found genes involved in auxin signalling in the M8 module: *EgrIAA27*, *EgrIAA19* and *EgrIAA33*.

However, we were able to identify several CW-related genes enclosed in modules of genes that are induced during the cold treatment and that could account for the cold-induced SCW remodelling.

First, we were able to identify genes belonging to the TRN controlling SCW deposition that were induced during cold treatment. We found early regulators of xylem differentiation, namely *ATHB8* and *REV/IFL*, but also NAC TFs belonging to the upstream regulators of SCW deposition, for example, *EgrNAC75* (orthlog of *AtVND7*, an early NAC master regulator involved in vessel specification) and *SND3*. We found also MYB TFs belonging to the bottom layers of the regulatory network controlling SCW deposition, such as *EgrMYB140* and *EgrMYB82*.

Then, even though the vast majority of CW biosynthetic genes were enclosed in modules M7 and M8 and therefore repressed after the beginning of the cold treatment, there were CW biosynthetic genes that were induced in this kinetics. Regarding cellulose biosynthesis, we retrieved sucrose synthases, hexokinases, phosphoglucomutases, UDP-glucose-pyrophosphorylases, involved in the UDP-glucose production and also cellulose synthases and cellulose-synthases like, involved in the polymerization of glucose to produce cellulose microfibrils.

Many hemicellulose biosynthetic genes are also induced in this co-expression network. Genes involved in hemicellulose backbone biosynthesis (UDP-xylose synthases) or addition or modification of side chains (*TBL* genes, *RWA* genes, *GXMT* genes) and hemicellulose modification (*GATL2*, *GATL10*) were found to be up-regulated by cold.

Even though most of the *bona fide* enzymes of the lignin pathway are found in M7 and M8 and thus only expressed at the beginning of the cold stress, some key biosynthetic lignin synthetic genes are induced during the cold treatment, namely *EgrF5H2* and *EgrCAD2*, catalysing the last step of monolignol biosynthesis. We also found laccases that were induced during the cold treatment, responsible for monolignol polymerization in the cell wall, enclosed mainly in module M9.

Finally, we also found genes involved in auxin signalling that were up-regulated by cold stress: *EgrIAA3.1*, *EgrIAA11*, *EgrIAA14*, *EgrIAA15.2*, *EgrIAA26.2*, *EgrIAA29*, *EgrIAA32, EgrARF10* and *EgrARF16.2*. Among them, *EgrIAA11* and *EgrIAA29* were previously found to be preferentially expressed in vascular tissues (Yu et al, 2015).

### 3.3. Wood related genes nested in CBF subnetwork are direct targets of CBF

Among the 4 344 cold-induced genes in the co-expression network, we found key members of cold signalling pathway, the *CBF* genes. 13 *CBF* genes out of the 17 reported in *E.grandis* genome were found nested in M1 module. They were directly linked to 1 257 DEGs of the global network (representing 15.2% of the network), belonging to 4 different modules (M1, M2, M10 and M11). Among these *CBF* genes, *EguCBFQ*, which was previously identified for its role in wood remodelling (Cao et al, 2020), was the most rapidly induced with the highest intensity (Figure S3). We hypothesized that *EguCBFQ* could be involved in cold-induced SCW remodelling. To decipher how EguCBFQ could remodel wood in response to cold, we used DAPseq to identify genes that were direct targets of this transcription factor.

We performed DAPseq experiment and obtained respectively 56 711 860, 30 531 040 and 33 406 070 raw reads for the three *Egu*CBFQ technical replicates (Figure 4A). After adaptors sequences removal (TrimGalore), *E.gunnii* genome mapping (BWA-MEM software), PCR duplicates elimination and filtering (threshold= 30), we obtained proper paired-end reads (9 953 068, 7 774 989 and 8 645 970 for each of the 3 CBF technical replicates (Figure 4A). Peaks calling with MACS2 generated between 24 493 and 29 578 peaks per technical replicate (Figure 4A). We then overlapped the peak sets from the 3 replicates to obtain a common peak set of 16 216 peaks. Analysis of motif enrichment with MEME-ChiP software revealed a significant enrichment in the known CBF-binding motif (CRT/DRE motif: CCGAC) within the common peak set (Figure 4B). To reduce the peak set and achieve a list of robust *Egu*CBFQ targets, we filtered the data according to (1) the presence of the CRT/DRE motif within the TF-bound sequence, (2) a DAPseq enrichment value above 5 and (3) a differential expression during the cold acclimation and freezing stress. Applying these criteria to the initial DAPseq peak set led to a list of 4 148 high-confidence targets (Figure S4). These 4 148 robust targets displayed a majority of peaks within the promoter region and after the 3’UTR (37% and 20% respectively), but also in the coding sequence (29%; Figure 4D).

**Figure 4:**
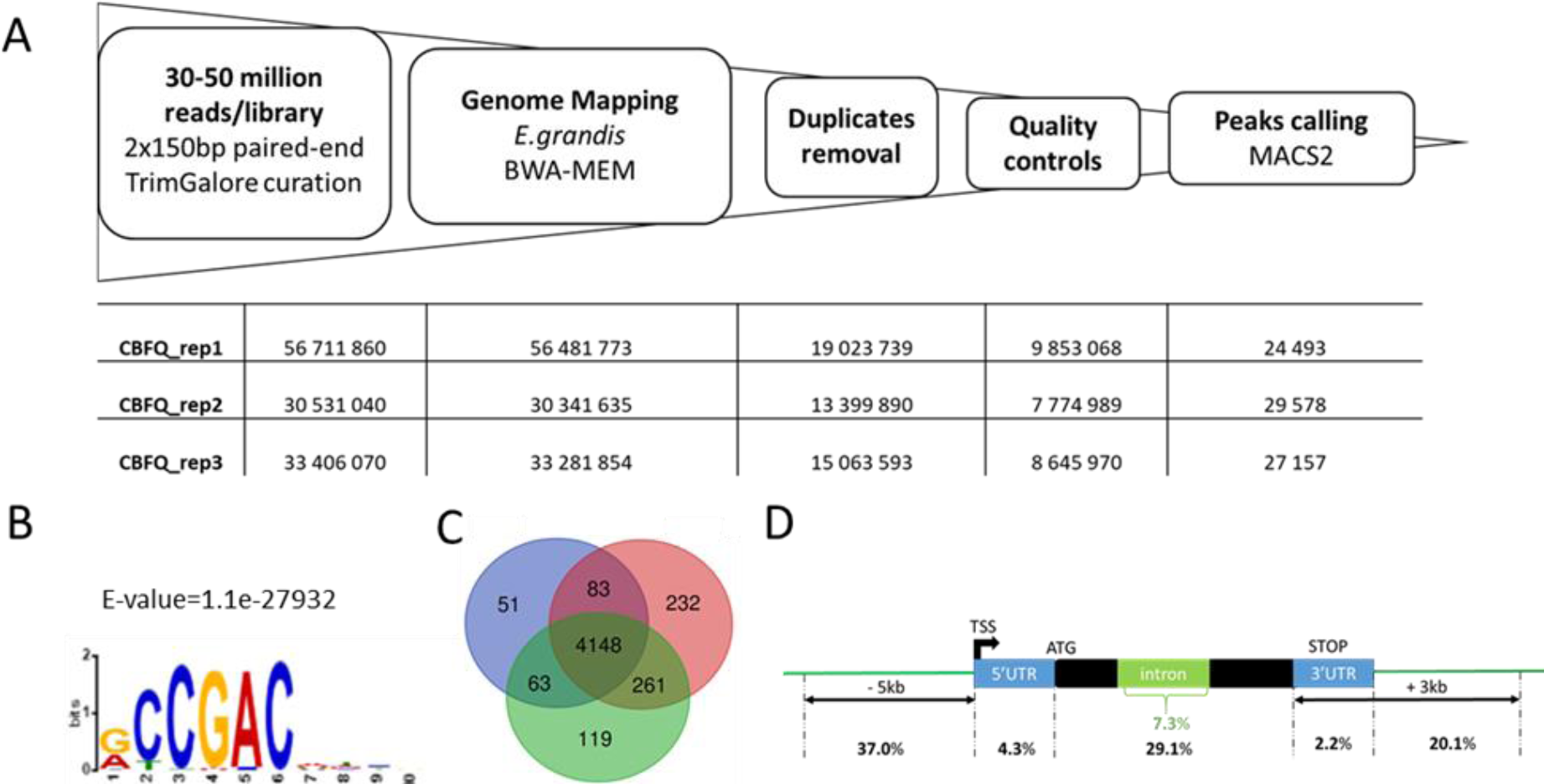
Genome-wide CBF cis-regulatory element detection by DAP sequencing. **A**: Analysis pipeline for curating raw reads from *Egu*CBFQ-DAPseq experiment. Number of reads are accounted for each technical replicate, at each analytical step. After peaks calling the number correspond to peaks number. **B**: Identification of conserved cis-elements motif in sequences of 500bp around the maximum of the peak performed using MEME-ChIP software (Machanick and Bailey, 2011) revealed the consensus CBF-binding motif in CBFQ-DAPseq peak set. **C:** Venn diagram of the genes identified in the three technical replicates of *Egu*CBFQ-DAP experiment after applying selection criteria (Figure S4) **D**: Repartition of the *Egu*CBFQ-DAP binding sites from the robust target list (4148 genes) along the gene structure.

We then investigated the functional categories of these 4 148 *Egu*CBFQ target genes using a Gene Ontology (GO) enrichment approach. 129 biological processes (GO categories) were significantly enriched in our list of DAPseq targets (Figure 5). We classified these categories in a hierarchized manner, from the most general (Level 1) to the most specialized (Level 10). We found a lot of level 10 categories related to abiotic stress response (10 GOs), biotic stress (3 GOs), biological regulation (4 GOs), transport (3 GOs), development (3 GOs) and especially plant-cell wall biogenesis (1 GO), signalling (1 GO), secondary metabolism (1 GO), primary metabolism (15 GOs), among which carbohydrate and polysaccharides metabolism (3 GOs), that could be related to cell wall formation. Besides, we found 1 level 10 GO category related to secondary metabolism and flavonoid biosynthesis pathway (Figure 6). Altogether, GO terms directly linked to cell wall biosynthesis or potentially related to cell wall metabolism encompass 264 DEGs: 125 of which are clustered within repressed modules and 139 are found within up-regulated modules.

**Figure 5:**
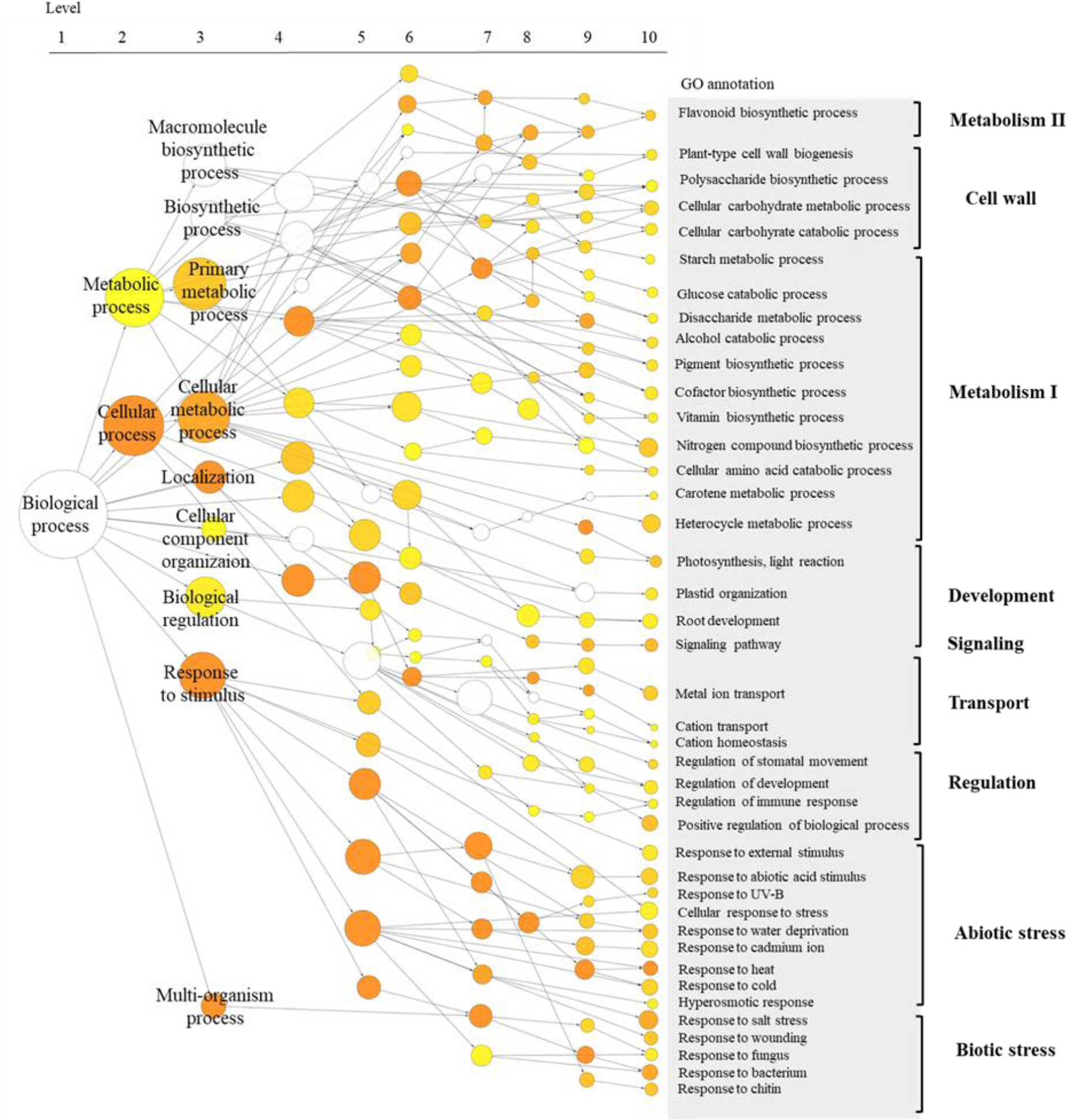
*Egu*CBFQ-DAPseq targets are enriched in stress and polysaccharides related GO terms. Gene Ontology (GO) enrichment analysis on the 4 148 *Egu*CBFQ-DAPseq targets. The network was created using BiNGO (Maere et al., 2005) with manual reorganization. A total of 129 significant GO terms (Pajdusted <0.01) were kept. Node size is proportional to the number of genes found in each GO category. Node colour is associated with statistical significance (from yellow to orange).

**Figure 6:**
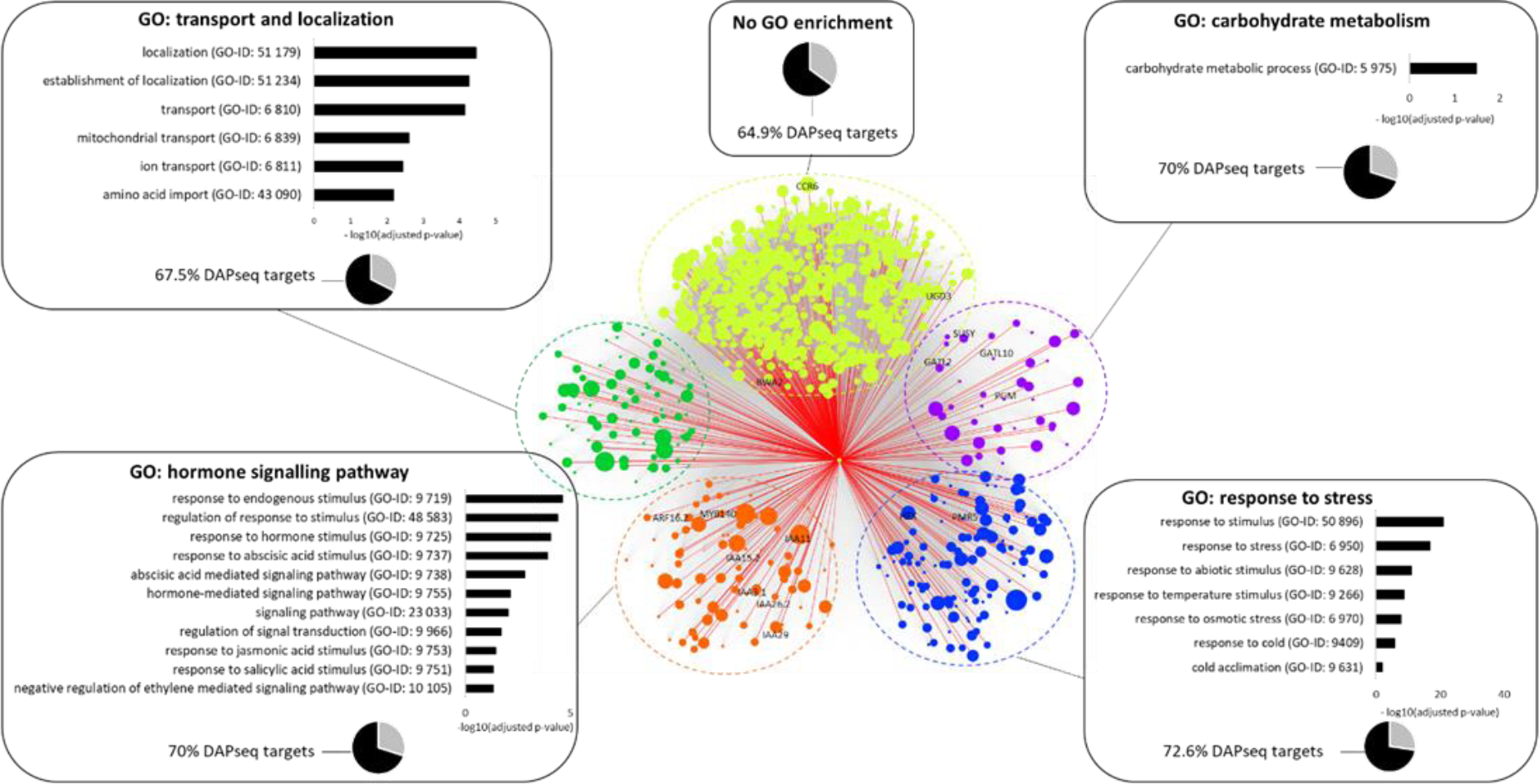
Identification of genes potentially involved in wood tissues remodelling within CBF subnetwork using co-expression and DAPseq. We extracted *Egu*CBFQ subnetwork (1105 direct neighbors, 515 620 edges) from global co-expression network (Fig. 3). Nodes are the 1105 direct neighbors of CBF extracted from the global network in Figure 3, linked by 515 620 correlations. Genes were grouped and colored according to Gene Ontology enrichment using BINGO, resulting in 4 GO-enriched modules and one module with no clear GO enrichment (based on orthology between *E.gunnii* and *A.thaliana*). 91 genes directly correlated with CBF had no close Arabidopsis ortholog and were discarded from analysis. Labels indicate well-characterized SCW-related or auxin-related genes. Pie charts represent the proportion of DAPseq targets among the module. Histograms display the p-value associated with each relevant GO term. Node size is proportional to DAPseq enrichment value. Grey edges represent direct correlations between genes and red edges link CBF with its DAPseq targets.

Besides, we found 1 level 10 GO category related to secondary metabolism and flavonoid biosynthesis pathway (Figure 6). Altogether, GO terms directly linked to cell wall biosynthesis or potentially related to cell wall metabolism encompass 264 DEGs: 125 of which are clustered within repressed modules and 139 are found within up-regulated modules.

To gain insight into the biological responses by CBF TFs, we extracted *Egu*CBFQ subnetwork (*i.e.* all the DEGs directly correlated with *Egu*CBFQ, with FDR-adjusted p-value >10^-6^) from the global co-expression network presented in Figure 3. We obtained a dense subnetwork of 1106 nodes linked by 515 620 correlations (Figure 6). Gene Ontology enrichment analysis on these 1106 *Egu*CBFQ highly correlated genes revealed 4 main GO categories, that were used to represent the subnetwork in Figure 6. These 4 main GO categories encompassed 312 genes of the subnetwork: 117 DEGs were associated to stress-related GO terms, 75 with intracellular transport and ion homeostasis, 80 with hormonal signalling pathways and 40 with carbohydrate metabolism. 91 DEGs highly correlated with *Egu*CBFQ had no *Arabidopsis* ortholog and were thus removed from analysis, leaving 701 genes with no significant GO enrichment.

Among these 1106 direct neighbours, 670 genes were also putative targets of the CBFQ TF, identified by DAPseq (representing 60,1% of the nodes in the *Egu*CBFQ subnetwork). Interestingly, we found genes involved in SCW formation that were DAPseq targets and co-expressed with *Egu*CBFQ. We detected one gene involved in cellulose biosynthesis (*EgrSUSY*, encoding a sucrose synthase), 3 genes involved in hemicellulose biosynthesis (*EgrUGD3*, *EgrRWA2*, *EgrGATL2*), 1 SCW regulator (*EgrMYB140*) and 4 *Aux/IAA* genes (*EgrIAA3.1*, *EgrIAA11*, *EgrIAA26.2* and *EgrIAA29*).

## 4. Discussion

Trees, as perennial long living organisms with long distance water transport, are seriously threatened by the ongoing climatic changes (Allen et al., 2010). It is therefore timely and of upmost importance to understand the mechanisms allowing their resistance to challenging environmental cues. One remarkable adaptive response of trees to stresses is wood plasticity occurring at the anatomical, chemical and physical levels, which is proposed to contribute to the functional adaptation of trees to environmental variations (Zinkgraf et al., 2017). However, the most important wood traits required to provide adaptation to a given stress have not been well-documented neither the molecular processes involved. Impact of cold on trees is supposed to remain a major issue in the future since the models of climate evolution predict the occurrence of frequent events of extreme temperature, like sudden frost events, accompanying the global warming. Cell wall changes induced by cold have been studied extensively in crop plants but only a handful of studies have investigated wood formation in suboptimal temperature conditions. In non-dormant tropical trees, like eucalyptus, cold signalling pathways can regulate xylem cell differentiation as demonstrated by previous studies from our group (Cao et al., 2020; Ployet et al., 2018).

Long term cold exposure has been shown to trigger SCW thickening and increased lignification, along with induction of SCW regulators and structural genes (Ployet et al., 2018). Overexpression of CBF TFs, which are key players of cold signalling pathway caused secondary xylem remodelling similar to what was observed in cold conditions, suggesting their involvement in cold-induced wood modifications. *CBF* overexpression triggered SCW thickening, increased lignification, smaller vessels and an up-regulation of genes belonging to the lignin *bona fide* biosynthetic pathway (Cao et al., 2020). However, if the works of Ployet et al, (2018) and Cao et al, (2020) represent landmark studies illustrating SCW remodelling in response to cold-induced CBF pathway in *Eucalyptus*, they used *a priori* approaches and focused on a subset of SCW related markers.

In the present study, we applied large scale approaches to analyse xylem transcriptome, global SCW chemotyping analysis and genome-wide detection of CBF direct targets in order to gain a comprehensive understanding about wood structure and composition remodelling mediated by CBF TFs in response to cold stress.

Taking advantage of a large phenotypic and transcriptomic data set, we showed that chilling and freezing temperatures trigger alterations in secondary xylem differentiation within cold-tolerant *Eucalyptus gundal* genotypes. We observed an increase of SCW thickness in developing xylem cells as well as selective lignification of cells surrounding vessels. We did not observe any change in global lignin content (data not shown). However, global analysis of SCW composition by FT-IR spectroscopy revealed changes in both lignin, hemicelluloses and cellulose shedding light on polysaccharides modifications occurring within the cell wall in response to cold exposure, suggesting a more global change than a simple increase in SCW thickness. Despite having pointed out the up-regulation of biosynthetic genes of these SCW compounds, previous studies had not explored further the modification of CW composition regarding polysaccharides. Besides alteration of SCW composition, chilling and freezing temperatures were associated with a decrease of cambial activity and a reduction of vessel size within the first developing xylem cell layers.

Transcriptomic analysis highlights that exposure to chilling and freezing temperatures triggers a deep transcriptomic reprogramming in xylem characterized by a high number of DEGs (8 236 DEGs out of 36 376 predicted genes representing 22.6% of *E.grandis* genome; Myburg et al., 2014). Modules of DEGs related to primary growth and metabolism, cell division and xylem development (M4 to M8), encompassing 3520 DEGs, are globally repressed during the cold stress, while stress-enriched genes are clearly-up regulated (2784 DEGs belonging to M1 and M2 modules). If cell-wall related modules (M7 and M8, encompassing 990 genes) are repressed after long-term cold exposure, it is worth noting that genes involved in SCW regulation, synthesis or deposition are found enclosed in other modules and induced during the cold treatment. Interestingly, modules enriched in GO terms related to cellular activity (M9 to M11, gathering 1560 DEGs), mainly transcriptional and translational activities, are induced, witnessing a high cellular activity going on during the cold treatment. This illustrates the strong regulation of cambial activity in this tropical species with continuous growth, that is yet influenced by environmental cues. These results suggest the existence of a transcriptional balance between growth and stress tolerance.

In this co-expression network, we found 13 out of 17 *CBF* genes, among which *EguCBFQ*, which has the highest and fastest induction of the *CBF* gene family. *EguCBFQ* is among the most induced genes in xylem during the freezing stress and is directly correlated to 1105 DEGs, suggesting its role as a regulatory hub during cold response in xylem. Given previous results showing that *EguCBFQ* overexpression triggers alterations of secondary xylem similar to those observed after long-term cold exposure (Cao et al., 2020; Ployet et al., 2018), we wanted to investigate how *EguCBFQ* can remodel wood formation by identifying the genes targeted by *Egu*CBFQ TF using DAPseq. We improved our list of target genes applying various criteria: (1) presence of the known CRT/DRE binding motif within the peak; (2) sufficient DAPseq enrichment value (>5) and (3) differential expression during cold treatment. We hypothesized that *Egu*CBFQ targets were more likely to share a similar expression pattern with *Egu*CBFQ and we took advantage of the co-expression network to select targets genes highly correlated with *Egu*CBFQ. Previous work from our lab used a machine-learning approach to refine DAPseq targets and revealed that co-expression data was the most important criterion (Takawira et al, in prep). Finally, the induction of the most promising *Egu*CBFQ targets was analysed in *CBF* overexpressing lines

Focusing on the direct role of *CBF* genes, we were able to identify 4148 putative direct targets of *Egu*CBFQ, corresponding to 3 541 unique *Arabidopsis* genes using DAPseq approach. As expected, given the known role of CBF transcription factors in cold response in plants in general and in *Eucalyptus* in particular, GO analysis of our putative target genes revealed a clear enrichment towards stress and cold response (363 DEGs were associated with the GO term “response to stress”). In herbaceous plants, CBF regulon has been extensively described through RNAseq, ChiPseq or DAPseq approaches. Park et al., (2015, 2018), have identified a CBF-core-regulon of 133 genes in *Arabidopsis*, corresponding to genes both up-regulated by cold and in *CBF*-overexpressing lines. 36.8% of their targets are found in our analysis, and our common set of targets are enriched in cold and stress response. O’Malley et al., (2016) have used DAPseq to identify direct targets of *AtCBF1* (closest ortholog of *EguCBFQ*) and it retrieved 883 genes in common with our DAPseq experiment (representing 25% of our target list). Finally, a recent study in *Arabidopsis* combining ChIPseq and RNAseq data has pointed out a CBF regulon composed of 146 genes, 40 of which are also in our analysis and are stress-related according to GO analysis (Song et al., 2021). Less data is available concerning CBF-regulon in woody plants, with to the best of our knowledge only one study addressing CBF targets in poplar (Li et al., 2017b). This study identified 2263 poplar CBF1 putative targets using a genome-wide strategy based on the detection of the conserved CBF-binding motif, with 217 *Arabidopsis* orthologs shared with our DAPseq target list, involved in stress response.

Even though these studies were conducted on different species (*Arabidopsis*, poplar, *Eucalyptus*), we found a significant overlap between our putative targets and previously described CBF regulons. Our common targets are enriched in GO terms related to stress and cold response and display an enrichment in the known CBF-binding motif (CRT/DRE motif). These results suggest a conservation of CBF signalling pathway and regulatory network between woody species and herbaceous plants. However, discrepancy between our target genes and previous CBF regulons could partially be due to the use of orthology to compare targets. For instance, 88.8% of *Eucalyptus* genes have an *Arabidopsis* ortholog, leaving a handful of genes deprived of *Arabidopsis* aliases, which we use in Gene Ontology analyses.

Finally, with our DAPseq analysis, we confirm *Egu*CBFQ role in stress response in *Eucalyptus*, with many targets being stress responsive, but we also uncover a link with CW metabolism and carbohydrates, with an enrichment in GO related to cell wall biogenesis (264 genes are linked to cell-wall related GO terms). Interestingly, when comparing our data with O’Malley et al., (2016), the list of overlapping targets is also enriched in cell-wall related terms, pointed out a new role of CBF TFs regarding cell wall remodelling in response to cold.

Our DAPseq results revealed promising candidates for the cold-induced SCW remodelling. *Egu*CBFQ DAPseq targets were enriched in GO terms related first to hormonal signalling pathways and second to the metabolism of sugars involved in SCW polymers structuration.

First, our results suggest that *Egu*CBFQ TF could bind to the promoters of *Aux/IAA* genes, *EgrIAA3.1*, *EgrIAA11, EgrIAA26*, *EgrIAA29* and *EgrIAA32* to induce their expression. We could hypothesize that CBF binds to *Aux/IAA* genes to repress auxin signalling and consequently influence xylem cell differentiation. This hypothesis is reinforced by previous studies unravelling a direct link between auxin signalling and *CBF* genes either in the model plant *Arabidopsis thaliana* or in perennial plants. Direct and physical link between *CBF* genes and members of the auxin signalling pathway was observed by Shani et al., (2017), with a ChIP experiment confirming the direct binding of *At*CBF1 on the promoters of two *Aux/IAA* genes (*AtIAA5* and *AtIAA19*). Further studies showed that *CBF* genes could regulate auxin signalling, triggering physiological effects. (Li et al., 2017a) dug up a strong link between CBF and auxin pathway thanks to a hormonal-focused transcriptomic analysis using *CBF* overexpressing lines, showing that more than half of the hormone-related DEGs were involved in auxin signalling, pointing out at the same time a clear repression of this pathway, accompanied with a reduction of IAA content. In *Malus domestica*, the overexpression of the peach *CBF1* gene (*Prunus persica*) impacts hormonal homeostasis, perturbing dormancy patterns (Artlip et al., 2019). They showed that genes involved in storage and inactivation of auxin were upregulated in transgenic lines exiting dormancy, while those for biosynthesis, uptake or signal transduction were generally downregulated. Targeting *Aux/IAA* genes could be a mean to trigger CW remodelling. Indeed, relationship between auxin signalling and wood formation has been established long before. The existence of a gradient of auxin along the cambium and the differentiating xylem, peaking inside the cambium has revealed auxin possible role in wood formation (Immanen et al., 2016; Tuominen et al., 1997) and triggered various studies investigating auxin role in xylem differentiation. In *Eucalyptus grandis*, two *Aux/IAA* genes have been already functionally characterized for their role in wood formation and secondary xylem patterning: *EgrIAA4* (Yu et al., 2015) and *EgrIAA20* (Yu et al., 2022). They were identified as preferentially expressed in vascular tissues in a genome-wide study of *Aux/IAA* genes in *E.grandis*, as were our candidate genes *EgrIAAA11* and *EgrIAA29* (Yu et al., 2015). *EgrIAA11* orthologs in *Arabidopsis thaliana*, *Populus trichocarpa* and in a *Populus tremula* hybrid are also preferentially expressed in xylem cells (Kalluri et al., 2007; Moyle et al., 2002; Yu et al., 2015) In *Populus tremula x Populus tremuloides*, *PttIAA5* (*EgrIAA11* closest ortholog) is strongly expressed in xylem cells harboring SCWs (Moyle et al., 2002). *EgrIAA29* ortholog in poplar (*PoptrIAA29*) is also expressed in xylem (Kalluri et al., 2007). By inducing the expression of *EgrIAA11* and *EgrIAA29*, CBF could indirectly contribute to repress auxin signalling, thus altering secondary xylem differentiation.

Second, besides targeting hormonal regulators of wood formation, our study pointed that *Egu*CBFQ could also directly bind the promoters of cell wall-related genes. SND3, which is a NAC TF master regulator of SCW deposition (Hussey, 2022), was identified as a putative *Egu*CBFQ target. In *Arabidopsis*, *SND3* is preferentially expressed in cells displaying SCW thickening and its dominant-repressor version triggers SCW reduction in fibres whereas its overexpression triggers SCW thickening (Zhong et al., 2008). In poplar, the fusion of SND3 with the activator domain VP16 induces ectopic deposition of SCW (Zhong et al., 2021). Given these data in both *Arabidopsis* and poplar, we could hypothesize that CBF could activate SND3 in response to cold, leading to the observed SCW thickening. Other MYB and NAC TFs are potential target of *Egu*CBFQ, including close orthologs of Arabidopis TFs involved in SCW TRN. Among them, we found *EgrMYB140*, whose closest orthologs in *Arabidopsis* are *AtMYB63*, well-described for its role in SCW lignification (Zhou et al., 2009) and *AtMYB15*, which is known for its role in cold response (Agarwal et al., 2006).

*Egu*CFBQ putative targets also encompassed genes involved in cell wall polysaccharides. A cell wall invertase (Eucgr.A02647) was among these targets. Cell wall invertases are responsible for the hydrolysis of sucrose into glucose and fructose (Tymowska-Lalanne and Kreis, 1998), and this source of glucose can be used for cellulose biosynthesis (Barratt et al., 2009; Mizrachi et al., 2012) but not exclusively. Its ortholog in *Arabidopsis* (AT3G52600) is a positive regulator of ovule initiation (Liao et al., 2020). As there is no clear link with secondary growth or xylem, its role in CBF-induced SCW remodelling needs to be further investigated.

We also found a galactosyltransferase (Eucgr.A01123) among *Egu*CBFQ putative direct targets. Its *Arabidopsis* ortholog (AT1G05170) was recently identified as involved in both primary and secondary cell wall biosynthesis (Nibbering et al., 2022). This gene encodes a CAGE2 protein (cellulose synthesis associated glycosyltransferase 2), localized in the Golgi and predicted to synthesize b-1,3-galactans on arabinogalactan proteins. The double mutant *cage1cage2* showed SCW defects such as collapsed xylem vessels and a reduction of SCW thickness in inflorescence stems, accompanied with a reduction in cellulose content and in SCW-associated CESA proteins (Nibbering et al., 2022). It appears as a promising candidate for the CBF-induced SCW thickening.

Finally, a pectin-lyase protein (Eucgr.J03054) was also identified among *Egu*CBFQ putative targets. Its *Arabidopsis* ortholog (AT1G80170) belongs to a group of xylem cells preferentially expressed genes (Ko et al., 2006). It is co-expressed with *AtERF38* in cells undergoing SCW thickening (Lasserre et al., 2008). Besides, a genome-wide study of pectin-lyase family in *Arabidopsis thaliana* shows that it’s induced in response to abiotic stresses, including cold (Cao, 2012). Pectin remodelling has been pointed out before in relation to cold response (reviewed in Le Gall et al, 2015).

Changes in wood structure and composition could contribute to plant tolerance towards freezing stress. The most striking cold-induced alteration of xylem anatomy is a reduction of vessel diameter. Reduction in vessel size has been previously associated with better tolerance to freeze/thaw embolism in trees, responding to the trade-off existing between xylem efficiency and xylem safety (Gleason et al., 2016). Besides, SCW thickening contributes to strengthen the cell wall, rendering it more resistant to cell collapse (Le Gall et al., 2015). Alterations of the biochemical composition of the cell wall could also contribute to mitigate plant tolerance to environmental cues. However, we still lack clear understanding of the role of CW modifications in the contribution towards freezing tolerance, and several studies reveal contrasting hypotheses. For example, in *Arabidopsis*, reduction of lignin content in SCW was associated with an increased freezing tolerance (Ji et al., 2015), whereas in *Rhododendron*, the accumulation of lignin and polysaccharides within the CW was associated to a better tolerance to freezing stress (Wei et al., 2006). In the *mur1* mutant in *Arabidopsis*, which displays less fucose in the cell wall, dimerization of pectin polymers is altered, reducing leaf freezing tolerance (Panter et al., 2019). In the same trend, overexpression of a pectin methylesterase inhibitor gene (*PMEI*), leading to alterations in pectin methylesterification patterns, triggered a reduction of freezing tolerance (Chen et al, 2018). Recently, Takahashi et al., (2021) showed that an *Arabidopsis* mutant in a *XTH* gene (xyloglucan endotransglucosylase/hydrolase), involved in xyloglucan remodelling, displayed a reduced freezing tolerance. Interestingly, we found that a *XTH* gene could also be targeted by CBF genes and constitute a promising candidate for understanding CW remodelling and freezing tolerance.

It is likely that CBF-induced modifications of wood structure and anatomy could contribute to better plant freezing tolerance but further functional characterization is needed to decipher precisely the molecular mechanisms behind it.

## 5. Acknowledgments

This work was supported by the Centre National pour la Recherche Scientifique (CNRS), the University Paul Sabatier Toulouse III (UPS, IDEX UNITI, project EuCoWood), the French Laboratory of Excellence project ‘TULIP’ (ANR-10-LABX-41; ANR-11-IDEX-0002-02) and the FR AIB (Fédération de Recherche Agrobiosciences, Interactions et Biodiversité, FR3450). The authors acknowledge F. Melun, J.Y. Fraysse, P. Alazard and Luc Harvengt (FCBA) for their help with the identification of field-grown eucalypts clones in Longages (France) and eucalypts culture in cold conditions, A. Gauthier and A. Desplat for their technical help during their Master internships, the Genotoul Bioinformatics Platform Toulouse Midi-Pyrenees for computing and storage resources and the TRI-Genotoul platform for microscopic analyses. The authors also acknowledge B. Potts, R. Vaillancourt and Jakob Butler (University of Tasmania), as well as W. Marande (CNRGV) for providing data from *E. gunnii* genome sequencing, assembly and annotation. R.P. and I.H.B. were supported by a PhD grant from the Ministère de l’Education Nationale, de l’Enseignement Supérieur et de la Recherche.

