## Supplementary information for "Cold stress involves CBF dependent regulatory pathway to remodel wood formation in *Eucalyptus gunnii* hybrids"

*Author for correspondence:

Mounet Fabien

### Supplementary data

#### Supplementary figures

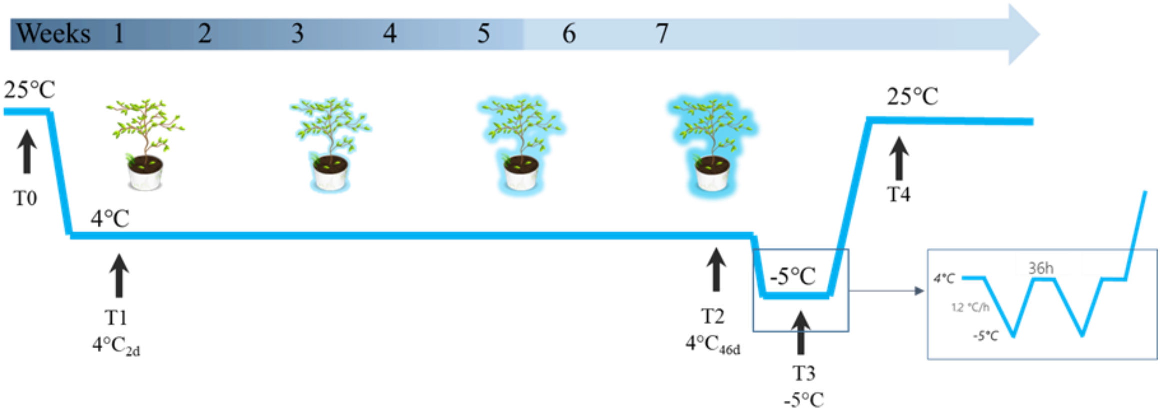


**Figure S1: Experimental set-up for the cold acclimation and freezing stress**

6-month old eucalypts (clonal copies of *E.gunnii* hybrids) provided by the FCBA (Pierroton, France) were submitted to a cold acclimation and a freezing stress. Plants were grown in a growth chamber at 25°C/22°C (16 h light/ 8h dark) and then submitted to a long term cold exposure at 4°C during 46 days, followed by a freezing stress at -5°C for 2 inconsecutive nights, before returning to 25°C. For the freezing stress, temperature decrease rate was 1.2°C/h. Stems were harvested on 18 plants per time point: at 25°C before the cold treatment (T0), 2 days after the beginning of the cold treatment (T1), 46 days after cold treatment (T2), after the freezing stress (T3) and 20 days after the return to 25°C (T4). For analyses of wood structure and composition, T0 and T1 samples are considered as “before cold” and T2 to T4 are considered as “after cold”.

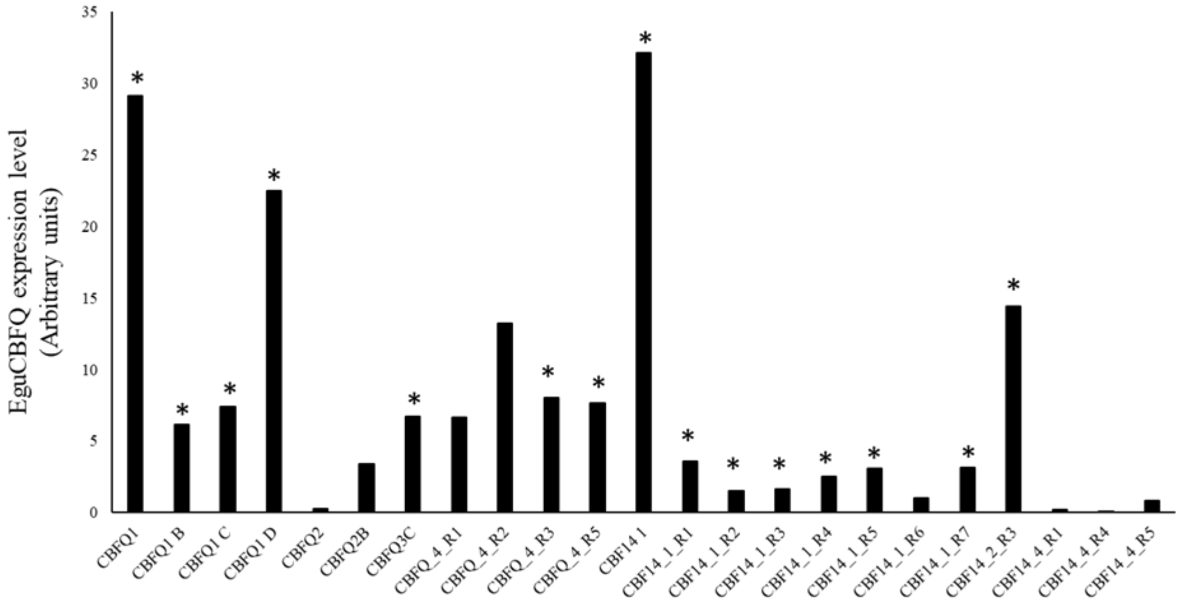


**Figure S2: Transgene expression level in *Egu*CBFQ transgenic lines.**

*EguCBFQ* expression was measured in control lines (empty vector). Asteriks represent the 15 root samples corresponding to 6 independent plants chosen for expression analysis. CBFQ_4_R1 and R2 were discarded according to root developing stage (absence of secondary growth)

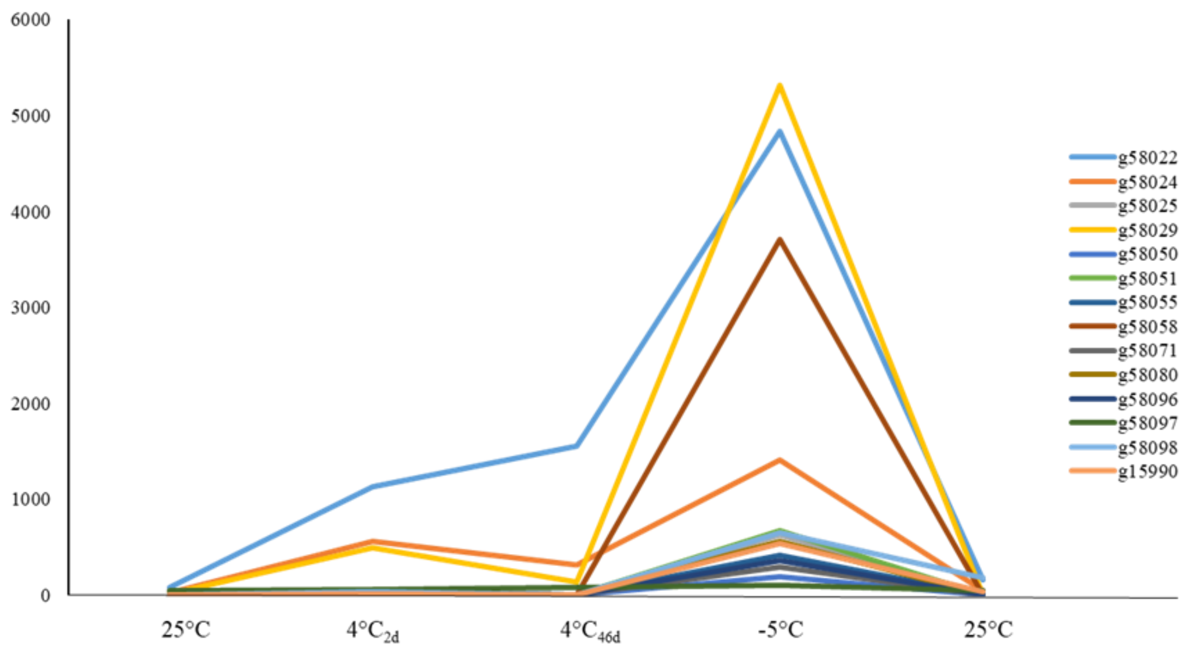


**Figure S3: *EguCBFQ* is the *CBF* gene most cold-induced during cold acclimation and freezing stress**

14 *EguCBF* genes were detected in the cold acclimation-freezing stress RNAseq. Mean expression value is calculated at each time point from 6 biological replicates and expression profiles are represented. Each *CBF* gene is named according to the temporary annotation of *E.gunnii* genome. *EguCBFQ* is g58022.

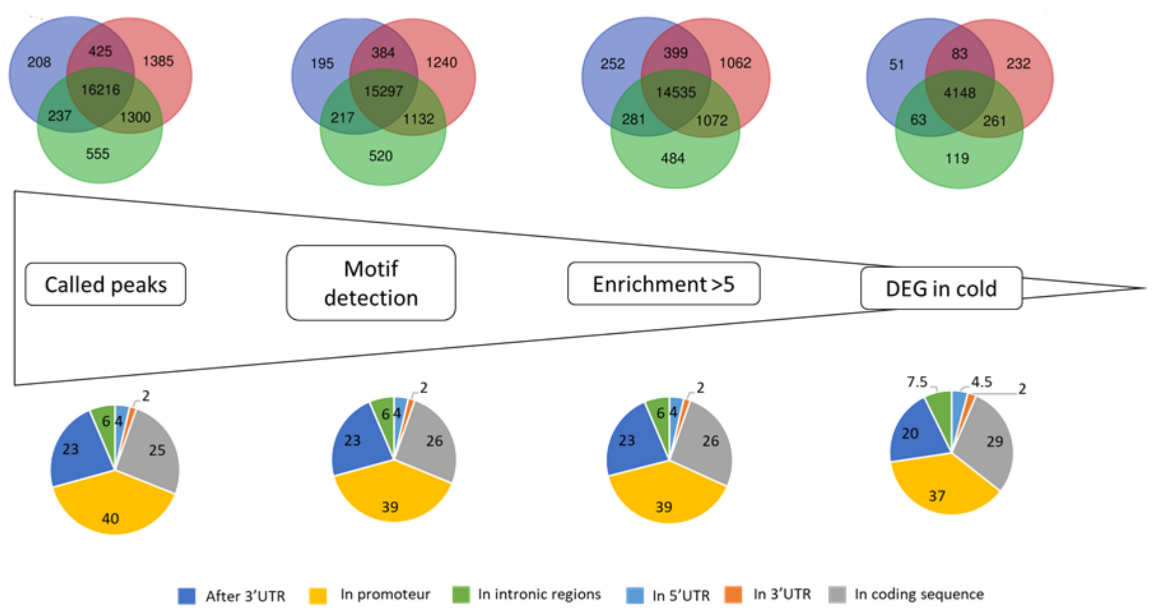

**Figure S4: Analysis pipeline to generate high confidence *Egu*CBFQ-DAPseq targets**

3 criteria were used to generate the robust target list: (1) presence of the CRT/DRE motif within the peaks, (2) DAPseq enrichment value >5, (3) gene differentially expressed during the cold and freezing kinetics. Up: Venn diagrams of the number of genes detected in the 3 *Egu*CBFQ-DAP technical replicates after the application of each selective criterion. Bottom: Corresponding localisation of the peaks along the gene structure. Values are mean percentage of the 3 technical replicates.

### Supplementary tables

**Table S1.** Primers used in this study.

| Accession | name | Annotation | primer F | primer R |
|---|---|---|---|---|
| **Cloning** | | | | |
| g58022 | EguCBFQ | CBFQ | CACCATGAACTCTTCCTCTTATATCTCCCA | TCACATGGAATAGCTCCATAATGAAGC |
| **Expression analysis** | | | | |
| g58022 | EguCBFQ | CBFQ | CCCATCCCAACTCCTTTAGCTTCG | TGGGACAGCAGCAACGAGAAAG |
| Eucgr.K03262 | MYB140 | MYB140 | AACAGAGCTCAAGCAGCGTGTC | TCCGGAGATTGCATCTTCAAGCG |
| Eucgr.H03171 | IAA3.1 | IAA3.1 | ACTCTGCTCCTTCTTCCAAAGCAC | GTTTCAGCCTCTGTCTTCTTCGTC |
| Eucgr.J02934 | IAA26.2 | IAA26.2 | TCTGAGATTTCTGTGCTCAGTGC | GCTCATTGCATGGCAGAATCCG |

| | | | | |
|---|---|---|---|---|
| Eucgr.C01734 | IAA29 | IAA29 | TCAAGTCCTGGAGGAAGGAGTTCG | TCTCGTAATGATCCTGGTGTGGTG |
| Eucgr.B03976 | RWA2 | RWA2 | GCATGGCTTGGGAAGATAACCC | TCCGGAATGCCTGACCTTAACC |
| Eucgr.K02506 | UGD3 | UGD3 | TCCCAGTTGGTGATTGAGAAGGC | GCTCCGATGCAACAAATCTTCACC |
| Eucgr.B03711 | HEX | HEX | AAATGGAGACAGATGCTGGAAGTC | AGCAGCTCTGCGGGTTACTATG |
| Eucgr.K01426 | IAA11 | IAA11 | TCAACGCACCACGTTCAAATAGG | TGAGGCACTTGTTGCTCGTACC |
| Eucgr.H01923 | GATL2 | GATL2 | GGAGAGCCTGCTACTTCAATACCG | AGTCACCTTCGCGCCATTTCTC |
| Eucgr.K03505 | SUSY | SUSY | GGCTGGAAACAGTCAGTGGAAC | GCAATTTCACCTGCTGCATCCTC |
| Eucgr.G00052 | CCR6 | CCR6 | TGTATGAGACACCGGCAGCATC | TGGTTTCTCCCGTAAACCTGTGG |
| Eucgr.E01031 | MYB68 | MYB68 | ACTACTGGAACTCGCACCTCAG | ACCGCCTCTTCTTCTTGTTGGC |
| Eucgr.A00969 | NAC10 | NAC10 | TCAGGAAGCGACTGAAAGACAGAG | TGGAAGTTTCCTCGAGCCATCC |
| Eucgr.F02864 | MYB82 | MYB82 | AGATGACGTCCCACTCATTGA | TGAACGGCAAATGCAGGTCCT |
| Eucgr.D01935 | KNAT7 | KNAT7 | CAAAGTGGCCATATCCAACTGAAG | GACCGATTGGGAATTGTTGTGC |
| Eucgr.B03684 | MYB31 | MYB31 | TTCGGGAGATTCGGAGGTGTT | TCGTTGCTGCTATTGTGTTCG |
| Eucgr.I02371 | F5H1 | F5H1 | ATTGAGACGAGGCATCGAACCG | CGCATGCAAAGCCAGCCATTTC |
| Eucgr.K00087 | 4CL2 | 4CL2 | TGGCTTTGACTGACGGAAGAAGC | ACTAGCCTTTCTGGCTGTTTGGC |
| Eucgr.B02853 | IAA32 | EgrIAA32 | GCGCATGCGGATCACAAGAAAG | TTGGTTGTTCGTCGGCTGATGC |
| Eucgr.B03636 | TBR | TBR | CCTATGGCCACTTTAGAGGAGGAC | TCTTCTTTGAATGGATCGGTCTCG |
| Eucgr.H03936 | UXS2 | UXS2 | TTCCAGTACGTTTCCGACCTGGTG | ATGGACCGACATGTTCGCCTTC |
| Eucgr.E03226 | SND3 | SND3 | TTAGAGGGAGAGAATGGGATCTGC | CCATCTTTGCTCACTCCTGGTAGC |
| Eucgr.E01499 | PMI | Plant invertase/pectin methylesterase inhibitor | TCTCCAGTAGTGGCAAAGACATCG | GGAACGTAAGCCGTTCGTAATACC |
| Eucgr.A02647 | CWI2 | cell wall invertase 2 | TGGGCTGGGATTCAGACGATTC | CCGCAGAGTTTCTAGCTCTTCAAC |
| Eucgr.F04341 | | NAC (No Apical Meristem) TF | | |
| Eucgr.D02103 | | myb-like transcription factor family protein | ACGTCATTTCAAGGACGCCAAC | CCCATTGGCACATCAACCGATTC |
| Eucgr.E01095 | NAC014 | NAC 014 | ACAGTCTGCTCCTACCAAGGTC | TGACCGTCTCTCTCTGGTTCATTG |
| Eucgr.I02215 | | Auxin efflux carrier family protein | TGCAACCCTTGCTGCTATATTGGG | TGAGCTTCCTCAGCCACGAAATG |
| Eucgr.F03681 | CSE1 | cellulose synthase like E1 | TCTGGGCTCCTAATTGCTTGCC | AACCATGGGCTCGTAACCTCTG |
| Eucgr.J03054 | | Pectin lyase-like superfamily protein | TTTGAGCCTTGTCCCAATCAGAC | CCTTCACATGGAGCATCATCACTG |
| Eucgr.H01338 | | Pectinacetylesterase family protein | GCAGATTCAGTTCCTGCAAGGC | TATCGGCGAACCATGTGTCCTG |
| Eucgr.C00176 | | xyloglucan endotransglycosylase 6 | TTGCATGGTGGATGTGGGTTGG | GCGGTTAGTGTTGCAACTTGGC |
| Eucgr.C01240 | CCR4 | CCR4 | TCGACATTGGCTTATCTCAAAGGG | CAGGAAATTAGTGTCGACGGATGC |
| Eucgr.B02574 | GATL9 | GATL9 | TCCGGGAGCACATTGATGATGG | ACACCGCCTATCAAAGAATCATCC |
| Eucgr.H01509 | TBL11 | TBL11 | TGTGTGACTTGCCCAGATTTGATG | ACCAACGAACACGACCCTTTGG |

| | | | | |
|---|---|---|---|---|
| Eucgr.F01647 | HEX | HEX | TTGTGAGGCGAGCTCTAGTGAG | TGCGGATAGATCTGGTGTCCTG |
| Eucgr.A02883 | PMI | Plant invertase/pectin methylesterase inhibitor | AGCCCTTCATACCGAGAGAAGC | TTCGCTTCTGAGGCTGCCATTC |
| Eucgr.K02102 | ARF | auxin-responsive family protein | ACGATGAAGGAGCAACATCTTCCC | TCGCACTAAACACACGGTAAACG |
| Eucgr.H05089 | | NAC (No Apical Meristem) TF | AAATGCGCTTCTCAGGCCATCG | ACAAAGCCATGTCTGGGAGCTG |
| Eucgr.A01123 | GST | Galactosyltransferase family protein | TGGAGCACATTGACGACAGGAG | TGGGCCTTCCATTCACAATCAG |
| Eucgr.F01093 | | NAC (No Apical Meristem) TF | TGCAAGAGCGGAAGCACTAAGC | TCCAATGAAGAGGAAGGCGAATGC |
| Eucgr.B00655 | UDP89B1 | UDP-glucosyl transferase 89B1 | AGCCTGACGGAGCTTCAATGTG | CAAATTGGACAGGCCATCAACGG |
| Eucgr.E01335 | GRT | galacturonosyltransferase 1 | GCATGGTGACTGAGTCTTCAGATG | CGAGCAAGCTTTGCAGGTGAATC |
| Eucgr.D01926 | UDPgal | UDP-galactose transporter 2 | CCTGATACAACGCAGATGGAGATG | AAGGAAAGGCAAGCCGACTACC |
| Eucgr.B02473 | EF1α | EF1α | ATGCGTCAGACTGTGGCTGTTG | TTGGTCACCTTGGCTCCACTTG |
| Eucgr.F02901 | IDH | IDH | AATCGACCTGCTTCGACCCTTC | TCGACCTTGATCTTCTCGAAACCC |
| Eucgr.B03031 | PP2A3 | PP2A3 | CGGAAGAACTGGGTGTGTTT | CACAGAGGGTCTCCAATGGT |

**Table S2. Most discriminant FT-IR wave numbers associated with CW compounds from literature data.** 64 wavenumbers related to CW compounds were identified from Sparse-PLS-DA loadings contribution to PC1 axis explaining samples separation (see Figure 2)

| | *WN (cm-1)* | *Assignment* | *Related CW compound* | *References* |
|---|---|---|---|---|
| *1* | 1730-1742 | C=O stretch in unconjugated ketones | lignin | Faix, 1991 |
| *2* | 1730-1742 | C=O stretch in glucuronic acid | hemicellulose | Chang et al, 2014 |
| *3* | 1720 | | phenolic ester | Alonso-Simon et al, 2011; Largo-Gosens et al, 2014 |

| | | | | |
|---|---|---|---|---|
| 4 | 1717-1737 | C=O strech, unconjugated keton, carboxyl and ester groups | lignin | Stark et al, 2016 |
| 5 | 1667-1670 | ring conjugated C=O stretch of coniferalhdehyde/ sinapaldehyde | lignin | Stark et al, 2016 |
| 6 | 1650-1675 | conjugated carbonyl | lignin | Sammons et al, 2013 |
| 7 | 1643-1645 | ring conjugated C=C stretch of coniferyl/sinapyl alcohol | lignin | Stark et al, 2016 |
| 8 | 1636-1638 | absorbed water | cellulose | Chang et al, 2014; Salim et al, 2021 |
| 9 | 1630 | phenolic ring | lignin | Alonso-Simon et al, 2011; Largo-Gosens et al, 2014 |
| 10 | 1611 | C=O: Aldehyde groups | lignin | Salim et al, 2021 |
| 11 | 1600-1630 | | de-esterified pectins | Alonso-Simon et al, 2011; Largo-Gosens et al, 2014 |
| 12 | 1596-1600 | aryl ring stretch symmetric | lignin | Stark et al, 2016 |
| 13 | 1595-1610 | aromatic ring | lignin | Popescu et al, 2007; Largo-Gosens et al, 2014 |
| 14 | 1588-1600 | aromatic skeletal | lignin | Faix, 1991 |
| 15 | 1530-1540 | | lignin | Largo-Gosens et al, 2014 |
| 16 | 1515 | phenolic ring | lignin | Alonso-Simon et al, 2011; Largo-Gosens et al, 2014 |
| 17 | 1510 | | lignin | Largo-Gosens et al, 2014 |
| 18 | 1506-1513 | aryl ring stretch asymmetric, C=C aromatic symmetrical stretching | lignin | Faix, 1991; Sammons et al, 2013; Stark et al, 2016; Salim et al, 2021 |
| 19 | 1464-1466 | C-H deformation asymmetric | lignin | Stark et al, 2016 |
| 20 | 1452-1462 | C-H deformation in CH3 and CH2 | lignin | Chang et al, 2014 |
| 21 | 1452-1462 | CH2 symmetric bending on xylose ring | hemicellulose | Chang et al, 2014 |

| | | | | |
|---|---|---|---|---|
| 22 | 1447 | Aromatic skeletal vibration combined with C-H in plane deformation | lignin | Salim et al, 2021 |
| 23 | 1365-1372 | C-H bending | cellulose | Chang et al, 2014 |
| 24 | 1330 | C-O stretch of S ring | lignin | Sammons et al, 2013; Largo-Gosens et al, 2014; Stark et al, 2016 |
| 25 | 1320 | | cellulose | Alonso-Simon et al, 2011; Largo-Gosens et al, 2014 |
| 26 | 1317 | C-O carbonyl | cellulose | Salim et al, 2021 |
| 27 | 1317 | xyloglucan | hemicellulose | Alonso-Simon et al, 2011; Largo-Gosens et al, 2014 |
| 28 | 1312-1316 | CH2 wagging | cellulose | Chang et al, 2014 |
| 29 | 1294 | C-O stretching of syringyl ring | lignin | Salim et al, 2021 |
| 30 | 1280 | C-H bending | cellulose | Liang and Marchessault, 1959 |
| 31 | 1270 | C-O stretch of G unit | lignin | Faix et al, 1991; Sammons et al, 2013; Largo-Gosens et al, 2014; Stark et al, 2016 |
| 32 | 1200-1205 | OH plane deformation | cellulose | Liang and Marchessault, 1959 |
| 33 | 1160 | C-O-C asymetric stretch | cellulose | Chang et al, 2014; Alonso-Simon et al, 2011; Largo-Gosens et al, 2014; Salim et al, 2021 |
| 34 | 1127 | aromatic CH | lignin | Stark et al, 2016 |
| 35 | 1125-1166 | CO stretch in ester group of HGS lignin | lignin | Sammons et al, 2013 |
| 36 | 1125 | aromatic CH deformation of G units | lignin | Sammons et al, 2013 |
| 37 | 1120 | xyloglucan | hemicellulose | Alonso-Simon et al, 2011; Largo-Gosens et al, 2014 |
| 38 | 1109 | | cellulose | Largo-Gosens et al, 2014 |
| 39 | 1104 | | pectin | Alonso-Simon et al, 2011; Largo-Gosens et al, 2014 |

| | | | | |
|---|---|---|---|---|
| 40 | 1100 | pectin acetylester | pectin | Largo-Gosens et al, 2014 |
| 41 | 1097 | | pectin | Alonso-Simon et al, 2011; Largo-Gosens et al, 2014 |
| 42 | 1082-1085 | C-O secondary alcohols | lignin | Stark et al, 2016 |
| 43 | 1078 | arabinogalactan, xyloglucan | hemicellulose | Alonso-Simon et al, 2011; Largo-Gosens et al, 2014 |
| 44 | 1060 | | cellulose | Alonso-Simon et al, 2011; Largo-Gosens et al, 2014 |
| 45 | 1047 | pectin acetylester | pectin | Largo-Gosens et al, 2014 |
| 46 | 1043 | arabinogalactan | hemicellulose | Largo-Gosens et al, 2014 |
| 47 | 1041 | xyloglucan | hemicellulose | Alonso-Simon et al, 2011; Largo-Gosens et al, 2014 |
| 48 | 1040 | | cellulose | Alonso-Simon et al, 2011; Largo-Gosens et al, 2014 |
| 49 | 1035-1050 | aromatic CH | lignin | Stark et al, 2016 |
| 50 | 1017 | pectin acetylester | pectin | Largo-Gosens et al, 2014 |
| 51 | 1017 | C-O: carbonyl | cellulose | Salim et al, 2021 |
| 52 | 1014 | | pectin | Alonso-Simon et al, 2011; Largo-Gosens et al, 2014 |
| 53 | 990 | | cellulose | Largo-Gosens et al, 2014 |
| 54 | 975 | | arabinose | Alonso-Simon et al, 2011 |
| 55 | 966-990 | CH-CH bending | lignin | Sammons et al, 2013 |
| 56 | 952 | | pectin | Alonso-Simon et al, 2011; Largo-Gosens et al, 2014 |
| 57 | 945 | | galactose | Alonso-Simon et al, 2011 |
| 58 | 915-930 | aromatic ring | lignin | Sammons et al, 2013 |

| | | | |
|---|---|---|---|
| 878 | CH deformation, aromatic ring | lignin | Stark et al, 2016 |
| 863 | CH deformation, aromatic ring | lignin | Stark et al, 2016 |
| 855 | CH bending of G units | lignin | Sammons et al, 2013 |
| 823 | CH deformation, aromatic ring | lignin | Stark et al, 2016 |
| 817-832 | CH bending of S units | lignin | Sammons et al, 2013 |
| 815-855 | aromatic ring G unit | lignin | Sammons et al, 2013 |
